# The complex admixture history and recent southern origins of Siberian populations

**DOI:** 10.1101/018770

**Authors:** Irina Pugach, Rostislav Matveev, Viktor Spitsyn, Sergey Makarov, Innokentiy Novgorodov, Vladimir Osakovsky, Mark Stoneking, Brigitte Pakendorf

**Author notes:** Corresponding authors (IP), (BP).

## Abstract

Although Siberia was inhabited by modern humans at an early stage, there is still debate over whether this area remained habitable during the extremely cold period of the Last Glacial Maximum or whether it was subsequently repopulated by peoples with a recent shared ancestry. Previous studies of the genetic history of Siberian populations were hampered by the extensive admixture that appears to have taken place among these populations, since commonly used methods assume a tree-like population history and at most single admixture events. We therefore developed a new method based on the covariance of ancestry components, which we validated with simulated data, in order to investigate this potentially complex admixture history and to distinguish the effects of shared ancestry from prehistoric migrations and contact. We furthermore adapted a previously devised method of admixture dating for use with multiple events of gene flow, and applied these methods to whole-genome genotype data from over 500 individuals belonging to 20 different Siberian ethnolinguistic groups. The results of these analyses indicate that there have indeed been multiple layers of admixture detectable in most of the Siberian populations, with considerable differences in the admixture histories of individual populations, and with the earliest events dated to not more than 4500 years ago. Furthermore, most of the populations of Siberia included here, even those settled far to the north, can be shown to have a southern origin. These results provide support for a recent population replacement in this region, with the northward expansions of different populations possibly being driven partly by the advent of pastoralism, especially reindeer domestication. These newly developed methods to analyse multiple admixture events should aid in the investigation of similarly complex population histories elsewhere.

**Author Summary:** We developed a new method that lets us disentangle complex admixture events and date the individual episodes of gene flow. We applied this method to a large dataset of genome-wide SNP genotypes from a linguistically diverse sample of populations covering the extent of Siberia. Our results demonstrate that these populations have been in intense contact with each other, with most of them carrying more than two, and up to six different ancestries. Nevertheless, as shown by the dates we estimate, these various admixture events have taken place over a relatively short time, with the oldest dating to not more than 4500 years ago. Our results emphasize the enormous impact that contact has had on the genetic history of these Siberian populations, and the methods developed here will be of use in the investigation of admixture events in populations around the world.

## Introduction

Siberia is an extensive geographical region of North Asia stretching from the Ural Mountains in the west to the Pacific Ocean in the east, and from the Arctic Ocean in the north to the Kazakh and Mongolian steppes in the south (Fig.1). This vast territory is inhabited by a relatively small number of indigenous peoples, with most populations numbering only in the hundreds or few thousands. These indigenous peoples speak a variety of languages belonging to the Turkic, Tungusic, Mongolic, Uralic, Yeniseic, Chukotko-Kamchatkan, and Eskimo-Aleut families as well as a few isolates. There is also variation in traditional subsistence patterns: the populations of southern Siberia are cattle and horse pastoralists; those of the Amur region focus mainly on fishing, hunting, and gathering; the peoples of central and northern Siberia frequently practise reindeer herding in addition to hunting; and along the coast of the Chukotka and Kamchatka peninsulas the Eskimos, Chukchi, and Koryaks are sea mammal hunters. This linguistic and cultural diversity suggests potentially different origins and historical trajectories of the Siberian peoples and warrants further investigation.

**Fig.1.**
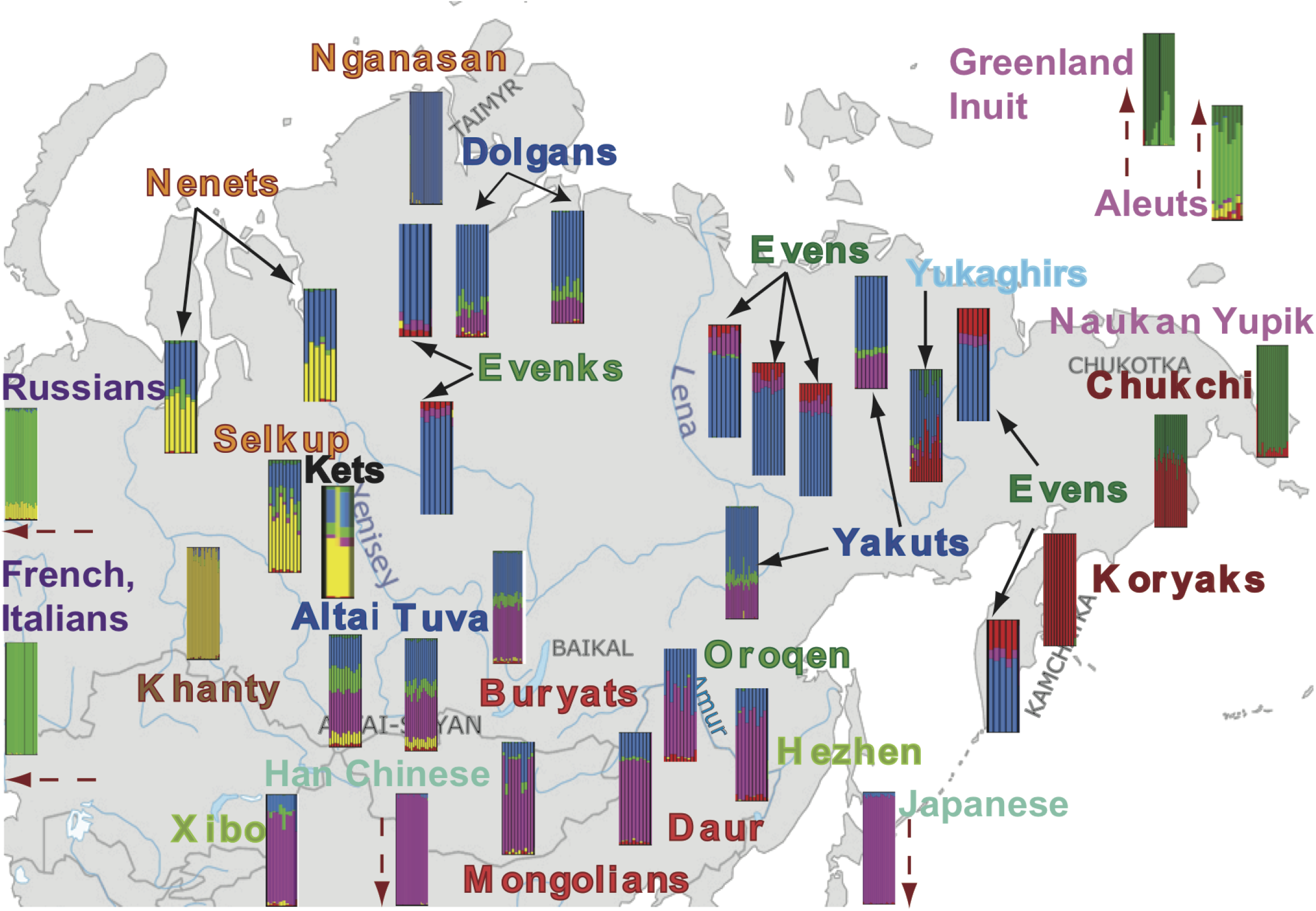
Admixture results for K=6 showing the approximate location of the populations included in this study. The names of the populations are coloured according to their linguistic affiliation as follows: red = Mongolic, blue = Turkic, dark green = North Tungusic, light green = South Tungusic (Hezhen) and Manchu (Xibo), brown = Ugric, orange = Samoyedic, black = Yenisseic, azure = Yukaghirs, maroon = Chukotko-Kamchatkan, pink = Eskimo-Aleut, purple = Indo-European, teal = Sino-Tibetan and Japonic. Where two subgroups are from the same geographic location, only one of the subgroups is shown (full results are presented in Fig.S1). Note that for reasons of space the location of the two distinct Yakut subgroups does not correspond to their true location. Each color indicates a different ancestry component referred to in the text as “(light) green” or European, “yellow” or Western Siberian, “blue” or Central Siberian, “pink” or Asian, “red” or Far Eastern, “dark green” or Eskimo.

The archaeological record attests to the ancient settlement of Siberia by modern humans. In particular, in the Altai-Sayan Mountains and the Lake Baikal region of South Siberia there is ample evidence of a long history of human occupation that highlights the important role South Siberia has played as a gateway into northeastern Asia and the New World. Anatomically modern humans were present in western and southern Siberia from as early as 46 kya [1–4]. In the Altai region they seem to have overlapped in time and might have co-existed with Neanderthals [5,6] and Denisovans [7,8]. The expansion of humans north was also rapid, and initial occupation of the Arctic environments is evident at more than 35 kya in the European part of the Russian Arctic [9]; by 27 kya humans are already in the Siberian northeast well above the Arctic circle [10,11], at a time when the Bering Land Bridge was still open [12].

An ongoing debate is centered on the degree to which human populations in Siberia were affected by the prolonged cold of the Last Glacial Maximum (LGM) 20-18 kya. Whereas some interpret the archaeological evidence as suggesting continued uninterrupted human occupation throughout the LGM [13–15], probably in protected locations with a milder microclimate, others argue that the extremely cold and arid conditions of the LGM led to a complete depopulation of southern and central Siberia [16–19]. It has been proposed that recolonization took place only when the climatic conditions improved after 18 kya, with new micro-blade stone flaking technologies appearing at this time in the archaeological record. Some researchers argue that these new technologies were brought to the area by immigrating people [17], while others view this change in technology as a gradual in-situ transition [20]. The genome sequence of the 24,000 year old individual from the Mal’ta site in southern Siberia reveals no genetic affinity between this Upper Paleolithic Siberian and modern human populations of southern and central Siberia, arguing for post-LGM population replacement, while intriguingly also revealing a genetic proximity to present-day Native Americans [21]. Evidence that the population replacement may have taken place at a much later stage (7,000-6,000 ya), however, comes from mtDNA analyses from two Neolithic cemeteries from Lake Baikal which are separated in time by a ∽800 year-long hiatus in settlement. While the pre-hiatus population shows affinities with western Eurasians, the post-hiatus population is genetically similar to modern-day populations of Southern and Central Siberia [22].

Despite Siberia’s evident importance for understanding the peopling of North Asia and the New World, there are few genetic studies which focus specifically on the history of the Siberian populations, and these are mainly confined to mtDNA and Y chromosome markers. Previous studies of ancient mtDNA and Y chromosome SNPs [23–25]and modern uniparental markers [26–31] highlighted the complexity of South Siberian history and the important role it has played in the peopling of North Asia and the Americas. In particular, the mtDNA pool of South Siberians is highly diverse [27,30,31], with different lineages tracing their ancestry to the Bronze and Iron Age periods [23–25,31] as well as to recent times [27]. Furthermore, the mtDNA haplogroups found in South Siberia are shared with other linguistically and culturally unrelated populations as far distant as northern and northeastern Siberia [28,30,32,33]. The populations of the Amur region and the Russian Far East, however, show distinct mtDNA lineages that testify to a separate history with partial links to the New World [33–36]. Only three recent studies were based on genome-wide data: one focusing on the population history of peoples of northeastern Siberia [37] and the other two focusing on selection involved in cold adaptation by Siberian populations [38] and in adaptation to the meat-rich diet in populations of the Chukotka peninsula [39]. These showed genetic affinities of the populations of Central Siberia with those of Southern rather than Northeastern Siberia, a signal of post-colonial European admixture in the populations of Central and Northeastern Siberia, and regionally and even population-specific signals of selection.

In this study, we compiled a comprehensive dataset of genome-wide SNP data from 20 Siberian and nine reference populations (Table S1) to investigate the relationships among the indigenous populations of Siberia. The assembled dataset includes new as well as previously published data [40–42] totalling 542 individuals, and covers nearly all of the indigenous language families and isolates represented in the region: Turkic, Mongolic, Tungusic, Uralic, Chukotko-Kamchatkan, Eskimo-Aleut, Yeniseic and the isolate Yukaghir languages (which might be distantly related to Uralic). The aim of our study is to provide for a better understanding of the genetic interactions between Siberian peoples, with a focus on disentangling shared ancestry from prehistoric migrations and contact. Commonly used methods to investigate admixture, such as TreeMix [43] and MixMapper [44], are inappropriate for the Siberian dataset, because the data strongly violate explicitly stated assumptions of these methods, i.e. instantaneous admixture at single time points and a tree-like population history [43,44]. Yet unraveling this complexity is essential for understanding the events and processes which shaped the human history of Siberia. We therefore had to rely on other methods and approaches, some of which were developed for this study. In particular, we introduce a new method for determining the order in which two or more admixture events occurred, which we refer to as an Admixture History Graph, and we modify a previous method for dating admixture [45] for use in more complex admixture scenarios. These new methods, when combined with other approaches (in particular, sharing of blocks of DNA that are identical by descent within and between populations), allow us to unravel some of the complex admixture history of Siberian populations.

## Results and Discussion

### The structure of genetic variation in Siberia

To understand the general patterns of relatedness between the samples in the dataset, we started with two widely-used exploratory tools: Principal Components Analysis (PCA) [45] and the model-based clustering algorithm ADMIXTURE [46]. The first principal axis (PC) is driven by differences between Europe and Asia, while the second PC differentiates the northeasternmost populations of the Russian Far East (Chukchi, Koryaks and Naukan Yupik) and the Inuit of Greenland (Fig.2). Although some populations fall where expected geographically and/or linguistically while still exhibiting potential signals of admixture, in several cases, the localization of populations in the PC-space is unexpected given their present-day area of settlement or their linguistic affiliation. Thus, the Mongolic populations (color-coded in red) fall close to Han Chinese and Japanese, except for the Buryats, who show closer affinities to the Turkic-speaking groups (color-coded in blue) than to other Mongolian populations. The Turkic-speaking groups of South Siberia (Altaians and Tuvans) and of Central and Northern Siberia (Yakuts and Dolgans, respectively) fall close together in the PC space, despite the large geographic distances that separate these populations. The Tungusic-speaking Evens and Evenks (color-coded in dark green), which were sampled all across central and eastern Siberia, cluster together and overlap with each other in the PC space. In contrast, the Oroqen, an ethnic minority group in northern China who are linguistically closely related to the Evenks [47], form a cline between the Tungusic peoples of Siberia and Han Chinese together with the other Tungusic-speaking minorities of northern China (Hezhen and Xibo; color-coded in light green). The Samoyedic-speaking Nganasan, who live on the Taimyr Peninsula in north Siberia (Fig.1) fall closer to the Evens and Evenks than to their linguistic relatives.

**Fig.2.**
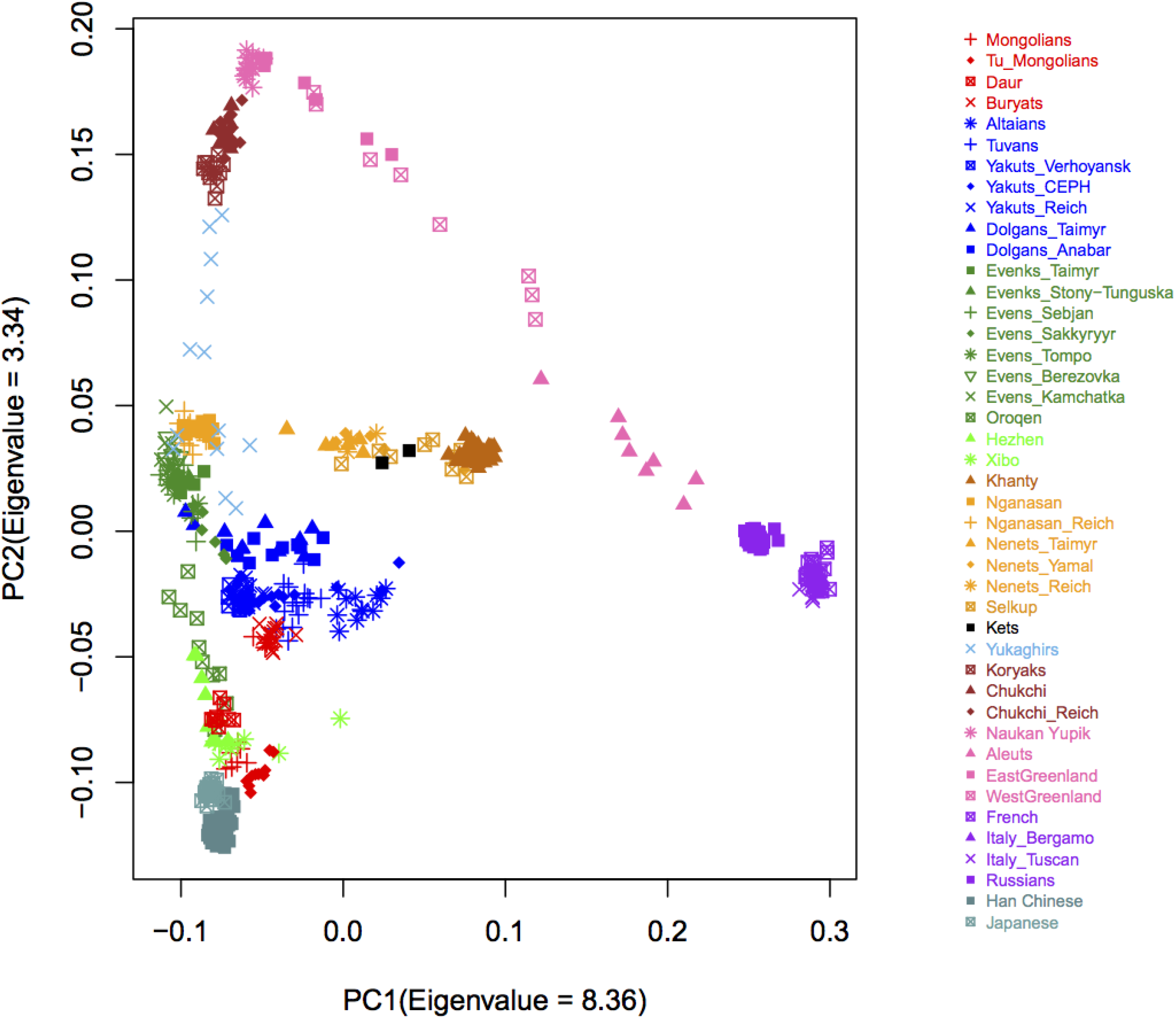
Results of the PC analysis showing the genetic structure captured by the first two principal components. Each coloured label represents an individual, and individuals are coloured according to their linguistic affiliation as described in the legend to Figure 1.

In order to investigate the patterns of potential gene flow, we estimated individual ancestry components using ADMIXTURE [46]. Results for K=6 are reported in Fig.1 (see Fig.S1 for the results obtained for K=3 through K=7). Although the lowest cross-validation error is obtained for K=7 (Fig.S2), at this value of K the Yakuts are differentiated from the other Siberian populations (Fig.S1), while at K=6 (for which the cross-validation error is also very low) Yakut ancestry is characterized in terms of components found in neighbouring groups, and hence these results are more valuable for understanding the prehistory of the Yakuts. Therefore, all AHG analyses and admixture dates reported here are based on the results for K=6. The six main components identified in this analysis can be roughly ascribed to Europe, East Asia, Western Siberia (present at highest frequency in the Khanty), Eskimo (present at highest frequency in the Naukan Yupik), Far East (Koryaks and Chukchi), and Central Siberia (found at highest frequency in the Nganasan). Strikingly, this analysis paints a very complex picture, revealing that most of the Siberians trace their ancestry to more than two, and sometimes to up to six, ancestral sources. These results suggest that admixture has been important in shaping the history of Siberia, but some of the observed variation seems to be of a clinal nature and follows a geographical gradient. For example, the Central Siberian (light blue) component is observed at highest frequency in the north, and decreases progressively southward. The opposite is true of the East Asian component (pink), while the Far Eastern component (red) seems to follow an east to west cline. While such clinal patterns could be indicative of admixture, they could also be explained by isolation by distance processes [48]. Hence, understanding and disentangling signals of common ancestry from signals of admixture is key to understanding the history of Siberia, and is the main focus of this study.

To better understand these patterns of relatedness and to elucidate to what extent the observed structure can be explained by recent local migrations, we inferred and analyzed chunks of DNA that were inherited without recombination by each pair of individuals from their most recent common ancestor, i.e. segments that are identical by descent (IBD) [49,50]. We expect individuals to share the highest number of, and longer, IBD segments with other individuals from the same population, and then (in the absence of long-range dispersals) with individuals from geographically close populations. In addition, we can use information on the amount of sharing within each population in comparison to other populations as indirect evidence of past population size changes, since the genomes of individuals in a population that has experienced a bottleneck have shallower genealogies, and hence are expected to share more IBD segments [51].

In terms of within-population IBD sharing, in general the individuals from the populations that now reside in the extreme North or the Russian Far East (e.g. the Naukan and Chukchi) share more IBD blocks with individuals from the same population than do individuals from populations with a more central location, such as the Altaians and the Tuvans (Fig.S3). This result is corroborated by the pattern of genome-wide linkage disequilibrium (LD), where the Koryaks and the Nganasan (populations from the Kamchatka and the Taimyr Peninsulas, respectively) exhibit much higher genome-wide LD than that observed for the Han Chinese or Europeans (Fig.S4A). This indicates that these populations living on the margins of the Siberian landmass likely experienced an extreme or several bottlenecks, possibly during successive migrations further north- and northeastwards.

With a few exceptions, the sharing of IBD blocks across Siberia is better explained by geographic proximity of the populations rather than by their linguistic affiliation (Fig.3 and Fig.S5). The most striking exceptions are the Altaians, Tuvans, and Mongolic populations, who share almost no IBD segments with any other population in the dataset, and the Evens, who share IBD segments even with geographically distant populations such as the Nganasan and Dolgans from the Taimyr or the Oroqen from North China. Such patterns of sharing indicate that although isolation by distance and recent local migration could explain most of the genetic variation in Siberia, they cannot account for all of the observed diversity. Rather, other demographic processes such as large-scale population dispersals and rapid expansion and/or gene flow (e.g. Evens) as well as language shifts (e.g. Buryats) are likely to have played a role in the history of some Siberian populations. Furthermore, the signal of recent interactions is completely absent in populations like the Altaians and Tuvans, and non IBD-based methods are needed to infer demographic events concerning these populations. One means of confirming prehistoric migrations is to identify the genetic traces of a population’s original geographical origin. It is has been previously shown that when PC analysis is applied to human genetic data from some geographic regions, such as Europe, the variation captured by the first two principal axes closely correlates with the geographic coordinates of the samples’ actual place of origin [52,53]. To further investigate the relationship between the distribution of genetic variation in our data and geography, we ran PC analysis on the inverse of the similarity matrix calculated from the number of shared IBD blocks between populations, as populations sharing more IBD blocks are more closely related genetically. The results are presented in Fig.4. For a better alignment with geography, the coordinate axes were rotated -45 degrees. The main axis of variation is oriented in a northeast – southwest direction, while the second axis separates east and west. Most of the genetic distances between the Siberian populations are smaller than expected based on their actual location on a geographic map (Fig.4). The most notable exception are the populations of the Taimyr, who are dispersed on the PC plot, even though they are settled in relatively close proximity to each other. In particular, the Nganasan appear to be much closer to the Tungusic-speaking Evenks and the Yukaghirs than to the neighboring Nenets (cf. [37]), even though the Nenets are not only geographically close to the Nganasan but also speak a related language. Similarly, the Mongolic-speaking Buryats cluster with the Turkic-speaking Altaians and Tuvans, and not with the Mongolic-speaking Mongolians and Daurs, although these are linguistically related and geographically less distant.

**Fig.3.**
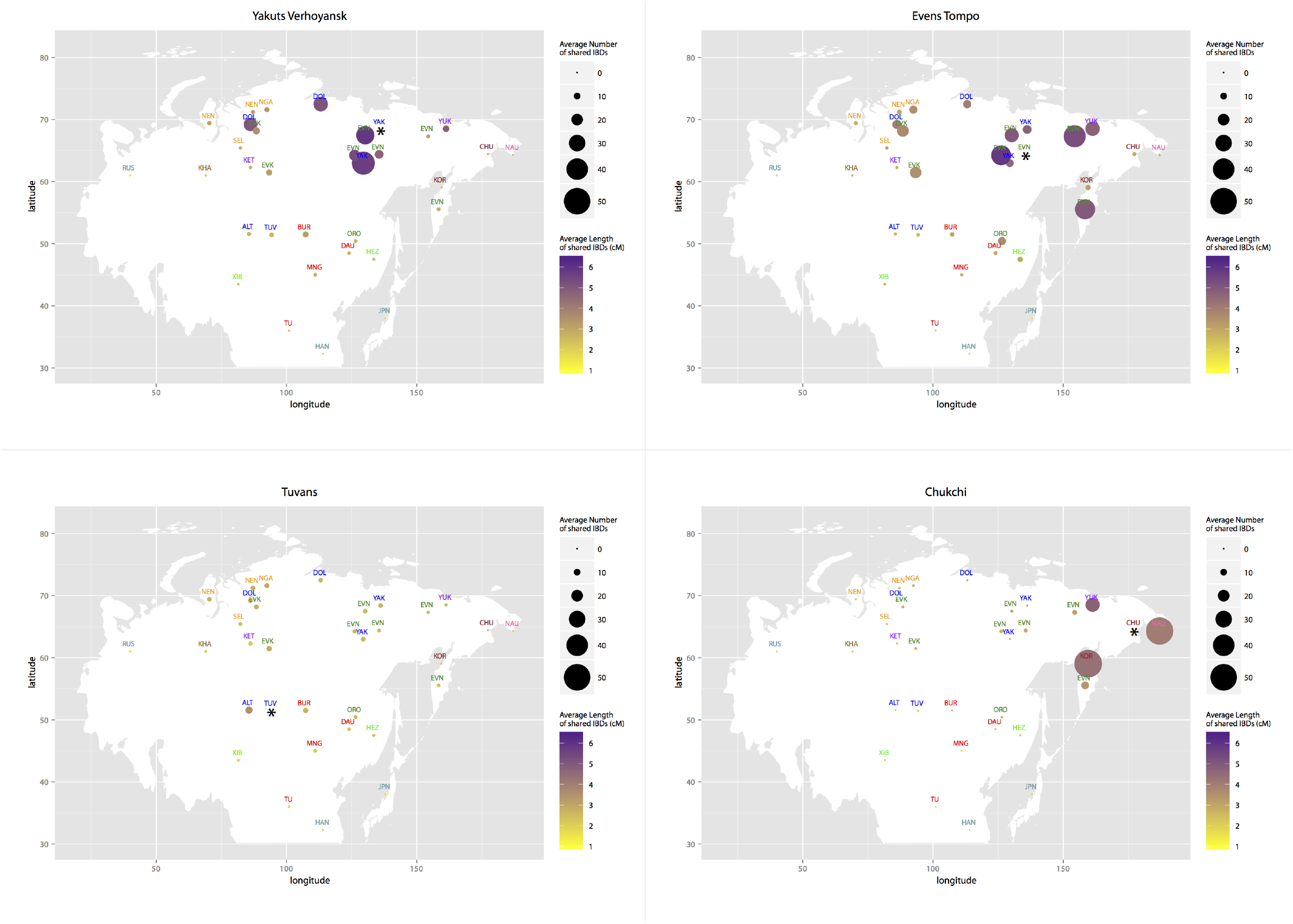
Recent relatedness as measured based on IBD blocks. Each data point represents the results for the comparison of the population marked with an asterisk to each of the other populations in the dataset. Data points are placed on the map according to the sampling location of each population (geographic coordinates are listed in Table S1). Population labels are abbreviated to the first 3 letters of the population name, except EVN = Even, EVK = Evenk, MNG = Mongolian, JPN = Japanese. Each label is color coded according to the population’s linguistic affiliation as described in the legend to Figure 1. The size of each circle is proportional to the mean number of IBD segments shared between the population marked with an asterisk and the population named in the label. The color intensity is proportional to the mean length of such shared IBD segments.

**Fig.4.**
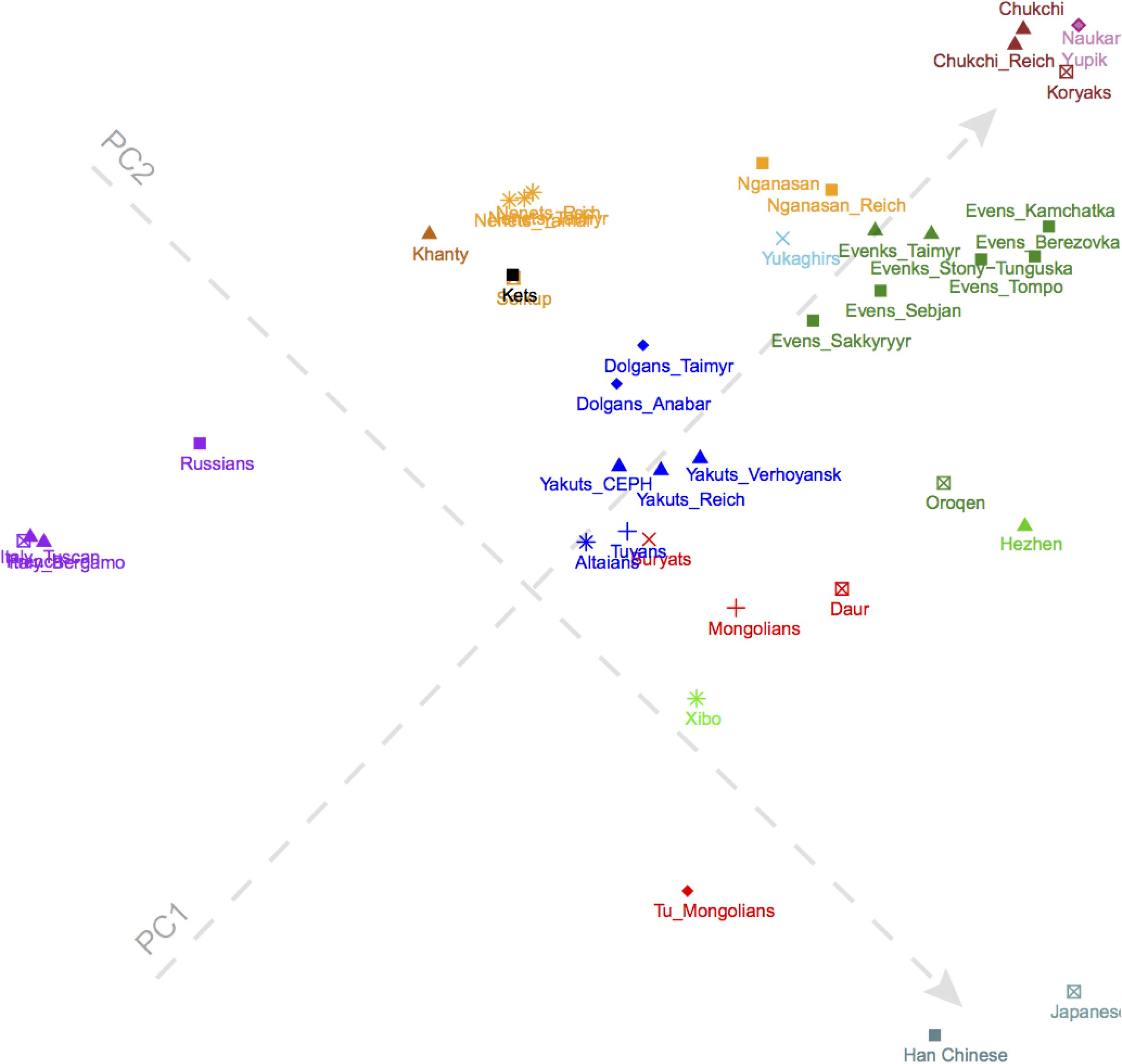
Results of the PC analysis based on the number of IBD blocks shared between populations. Each coloured label represents a population, and colour coding is according to linguistic affiliation, as described in the legend to Figure 1. The PC axes are rotated for a better alignment with the geographic map of Siberia.

To better understand how genetic distances between populations are influenced by geography and to what extent the genetic variation in Siberia might be explained by isolation by distance, we used multiple regression of genetic distances on geographic distances [54,55]. Historically, rivers played an important role in the movement of peoples across Siberia. Today, as in the past, when they freeze over in winter these rivers serve as “ice highways”, efficiently connecting far-away settlements and towns. Rivers therefore were also included into our regression model as an additional explanatory variable, as described in the Materials & Methods. This analysis reveals that geographic distance alone explains 48% of the genetic variance (p-value < 0.001). Adding rivers as facilitators of gene flow improves the fit of the observed genetic distances to geography, increasing the Rsq by 0.02 (p-value < 0.001). We then jackknifed over populations by sequentially removing each population and refitting the regression [54]. The largest improvement to the fit was achieved by removing the Aleuts, Han Chinese and Japanese samples, but still only 58% of the variation in genetic distances between populations is explained by geography (p-value < 0.001). The general conclusion from this analysis is that although geography explains a large part of the genetic variance, it does not explain all of the genetic relationships in Siberia. This further demonstrates that long-distance migration and admixture are likely to have played a substantial role in the prehistory of Siberian populations.

In summary, our exploratory analyses demonstrate that the prehistory of Siberian populations has been complex, with large-scale dispersals, local effects of geographical proximity, and variable degrees of admixture playing a role in structuring the genetic variation of these populations. These analyses also demonstrate that individual populations have had different historical trajectories, with the populations of the periphery being notably distinct from the others. In the following sections we therefore analyze signals of admixture and discuss the genetic prehistory of populations grouped either by their present-day geographical location or by their linguistic affiliation.

### South Siberia (Altaians, Tuvans, and Buryats)

The populations of southern Siberia are mainly pastoralists who speak Turkic (Altaians and Tuvans) or Mongolic languages (Buryats). Evidence for the use of domesticated animals (cattle, sheep, goats and horses) in this region goes back to the Neolithic [56–59]. The results of the ADMIXTURE analysis for the populations of South Siberia are striking (Fig.1) and reveal the important role played by the Sayan and Altai Mountains in the human history of Siberia: all of the ancestry components found across Siberia are represented here (cf. [38]), although two (Far East and Eskimo) are present in too low proportions to permit further analysis. This is consistent with South Siberia’s rich archaeological record with a wealth of sites representing different cultures, either contemporaneous or following in quick succession, and attests to its importance at the crossroads of various migration routes [60–62], which resulted in an amalgamation of different ancestries.

The ancestry composition in Altaians, Tuvans, and Buryats is almost the same (Fig.5A), even though the proportions of the different ancestry components vary slightly among the populations. The Altaians have a higher amount of the European component (21% vs ∽10% in Tuvans and Buryats) and a lower amount of the Central Siberian ancestry component (27% vs 37% in Tuvans and Buryats). The Western Siberian ancestry component occurs at similar frequencies in Altaians and Tuvans (10% and 8%, respectively), while it is almost absent in Buryats (2%). In contrast, the Asian ancestry component is higher in Buryats (47% vs ∽38% in Altaians and Tuvans).

**Fig. 5.**
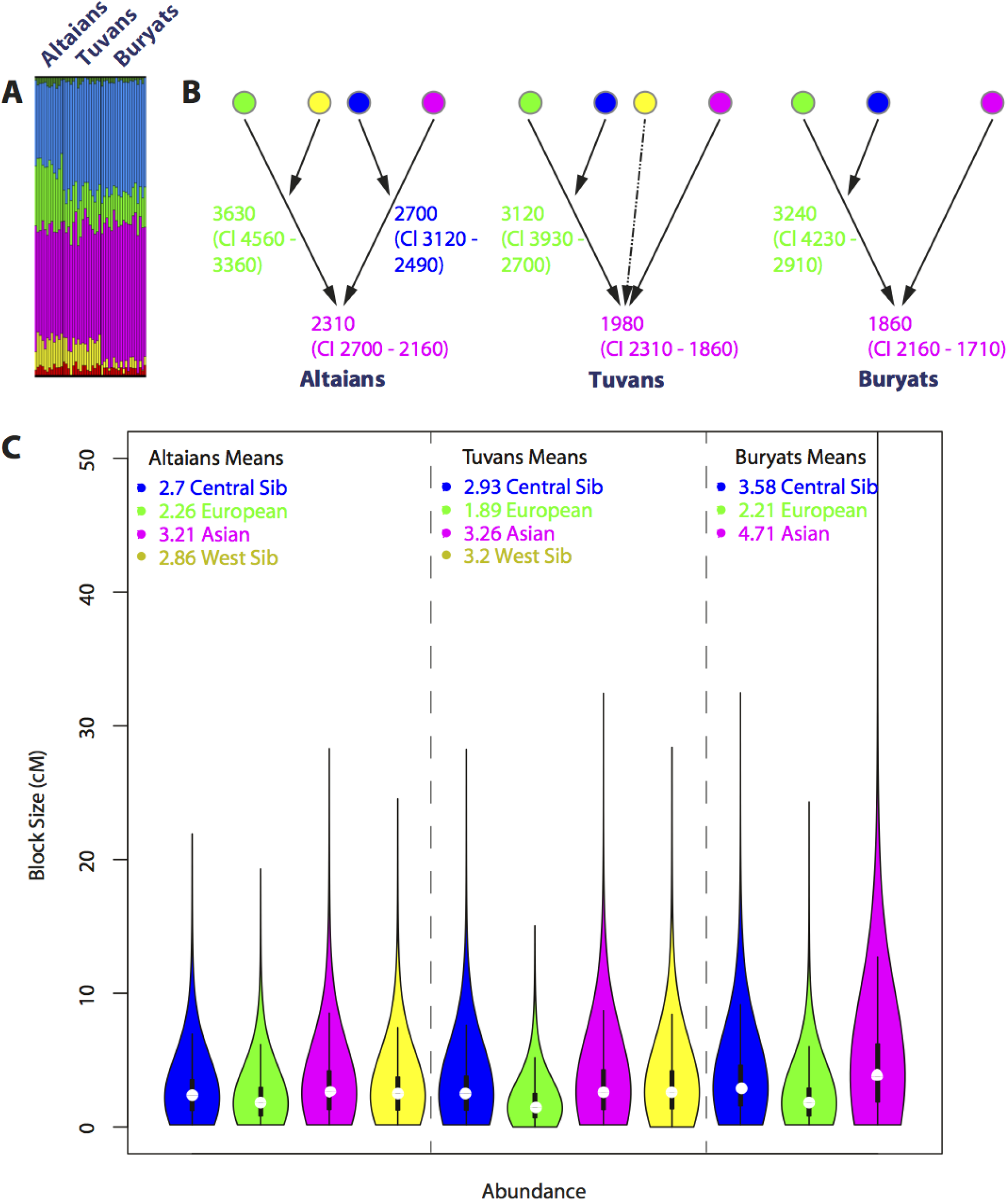
Admixture profiles for populations of South Siberia: Altaians, Tuvans, and Buryats. (A) An excerpt from the plot summarizing results of the ADMIXTURE analysis for the Altaians, Tuvans and Buryats at K=6. The full panel is shown in Figure S1. (B) Admixture history graphs and admixture dates inferred for each population and each admixture episode. Proxy parental populations for the different ancestral components (represented as circles) were as follows: European (green) = Italians, Western Siberian (yellow) = Khanty, Central Siberian (blue) = Nganasan, Asian (pink) = Han Chinese. (C) Cumulative distribution of all ancestry blocks. For each population the plot captures the total abundance of blocks of each ancestry (x axis) of different genetic lengths in cM (y axis); the average width of the blocks of each ancestry and variance around the mean are also shown.

Despite the overall similarity of the ADMIXTURE results, the AHG analysis infers different admixture graphs (and hence, admixture histories) for the Altaians as compared to Tuvans and Buryats (Fig.5B). However, it should be noted that there is some uncertainty in the reconstructed sequence of admixture events involving more than three ancestral populations when sample sizes are small and levels of admixture low (see TextS1). As both issues are relevant for South Siberia, we performed further tests to validate the different inferred admixture sequences we obtain for the South Siberian populations (see TextS1 for details).

The configuration most supported by the AHG calculation (Table S2) is shown in Fig.5B. As can be seen, the admixture history of Altaians differs from that of Tuvans and Buryats (Fig.5B and TableS2) in that the first admixture event in Altaians involved Western Siberian and European ancestries, whereas for Tuvans and Buryats the first event involved European and Central Siberian ancestries. The estimated dates of the admixture events, however, are roughly comparable, with the first event taking place ∽3,000-3,500 years BP and the most recent event ∽2,000 years BP. Since the average width of blocks of Asian ancestry and the variance around the mean are larger in Buryats than in Altaians and Tuvans (Fig.5C), it is likely that the Buryats experienced additional gene flow from a source of mainly Asian ancestry. This would explain why the signal of Asian admixture in the Buryats appears to be younger than the same signal in Altaians and Tuvans. The admixture history graphs and the dates of the admixture events inferred here for the South Siberian populations are consistent with the archaeological record (see TextS2 for details).

To summarize, although we cannot rule out the possibility that the admixture history in the three South Siberian populations analysed here was the same, the evidence instead suggests different scenarios for Altaians and Tuvans/Buryats. Since the European ancestry component is involved in the earliest admixture events in all three populations and the average length of European IBD tracts across the South Siberian populations is the smallest, this component cannot reflect post-colonial Russian admixture, in contrast to previous findings [38]. Instead, it appears to reflect the oldest population substrate in South Siberia (Fig.5B). The Western Siberian component first appeared in the Altai region, and then spread east to what is now Tuva and Buryatia (where it is seen at low frequency). The opposite pattern is observed for the Central Siberian component, with the older admixture dates inferred in Buryats and Tuvans and a later date in Altaians, implying that this ancestral component spread from east to west (Fig.5B).

The particular demographic history of the South Siberian populations is further underlined by the LD analysis (Fig.S4B): the short-range LD is lower in Southern Siberia than in the Han Chinese, while the long-range LD is higher than in the Chinese. Since long-range LD is informative about recent demography, while short-range LD carries information about events that occurred further back in the past [63,64], this pattern suggests that the stronger historical bottleneck that seems to have been experienced by Asians as compared to Europeans (e.g. [65,66]) is not observed in the populations of South Siberia. The ancestors of the South Siberian populations would therefore have experienced a weaker bottleneck than Asians, but a stronger bottleneck than Europeans. A possible reason for this might lie in the relatively hospitable environment of this region, which carried substantial population sizes even during the Last Glacial Maximum [13–15,19]. The distribution of the genome-wide LD reverses at larger genomic distances, where the level of LD in Europeans is the same as in Asians, while the values for Southern Siberia are elevated. It is known that long-range LD can be affected by either bottlenecks or migrations [67–70]. However, the results of the IBD segment analysis suggest that like Europeans and Han Chinese, but unlike most other Siberian populations (e.g. Yakuts, Nganasan, Koryaks), all Southern Siberians share relatively few IBD segments within the population (Fig.S3), a pattern commonly seen in non-isolated populations [50,71]. The higher long-range LD in Altaians, Tuvans and Buryats, in conjunction with high Ne (as suggested by their low genome-wide LD), is therefore consistent with these populations having experienced more recent gene flow, as is also shown by the admixture analyses.

It is furthermore notable that the Buryats, who speak a Mongolic language closely related to Khalkha Mongolian, genetically do not pattern with the Mongolians, but with the South Siberian Turkic groups in all respects. This is consistent with the prevailing view that Buryats are the descendants of indigenous populations from Lake Baikal who shifted to their current Mongolic language [72–74].

### Yakuts and Dolgans

Although they live in central and northern Siberia, the Yakuts and Dolgans speak Turkic languages related to those spoken in the south. The Yakuts practice animal husbandry like the South Siberian Turkic-speakers, whereas the Dolgans are nomadic reindeer breeders and hunters like their neighbours, the Tungusic-speaking Evenks [75–77]. Notwithstanding the great geographic distances that separate the Yakuts and Dolgans from their linguistic relatives, their genetic relationship with the South Siberian populations emerges clearly in both the PC and ADMIXTURE analyses. In the PC analysis, the Turkic-speaking populations are not separated along PC1-PC2, and a distinction between the southern (Altaians, Tuvans) and northern (Yakuts, Dolgans) Turkic-speaking groups is first observable only at PC3 and PC4 (Fig.S6). It has been shown before that for spatially structured data the first two PCs accurately capture geography, and genetic information can thus be used to infer geographic origin [52,53]. Such a genetic trace of pre-migration origins can be discerned for the Yakuts: their position in the PC analysis would place their geographic origin to the east of Tuva and north of Buryatia; this is in good accordance with their purported origins around Lake Baikal [75,78].

The hypothesis of a southern origin is also indirectly supported by the results of the IBD analysis. The amount of the IBD segments shared within the Yakuts is relatively high in comparison to other populations (Fig.S3); for instance, they share twice as many segments within the population as the Buryats, even though the census sizes of Buryats and Yakuts are comparable (Buryats – 461,389 individuals, Yakuts – 478,085 individuals; Russian 2010 census). Furthermore, the slow decay in the frequency of small IBD segments (2-10 cM) in comparison to longer segments in the Yakuts (Fig.S3) also suggests a strong founder event [79]. Although the Yakuts currently constitute one of the most numerous indigenous ethnic groups in Siberia, this pattern of sharing of IBD segments is consistent with a recent expansion from a small founding population, which is also supported by uniparental data [32,80,81] and by the relatively high genome-wide LD in the Yakuts (Fig.S4B).

In terms of sharing with other populations, the Yakuts share more and the longest IBD blocks with the Dolgans, who are thought to be partially descended from Yakuts ([82] and references therein), but also with the neighbouring Even groups (Sakkyryyr, Sebjan, Tompo), Buryats, and Evenks (Fig 4 and Fig.S5). The sharing of IBD blocks with Buryats is consistent with the idea that the Yakut ancestors once lived in close proximity to the ancestors of modern-day Buryats. The Dolgans share a large amount of long IBD segments with the Yakuts, Evenks, Nganasan and Evens (Fig.S5). While the Yakuts and Evenks are thought to have contributed genetically to the modern-day Dolgans, the affinity towards the Nganasan and Evens is probably indirect, and is best explained by these populations having themselves shared ancestry with the Evenks, although direct admixture with Nganasan cannot be excluded.

The ADMIXTURE results (Fig.1) suggest three sources of ancestry for the Yakuts and the Dolgans: Central Siberian (blue) (59% in the Yakuts, 67% in the Dolgans), Asian (pink) (30% in the Yakuts, 18% in the Dolgans), and European (green) (10% in the Yakuts, 11% in the Dolgans). In contrast to their South Siberian Turkic-speaking relatives, the Yakuts and Dolgans lack the Western Siberian component (yellow).

Analysis of the distribution of the widths of blocks of different ancestries reveals that the blocks of Central Siberian ancestry in Yakuts and Dolgans become progressively wider to the north and west (Fig.S7B). This would be consistent with additional recent gene flow from a population of mainly Central Siberian ancestry. In fact, the widest blocks and the highest variance around the mean are observed in the Taimyr Dolgans, the population with close genetic ties to the Evenks, who in turn have a high proportion of the Central Siberian ancestral component. All the inferred admixture dates given in Fig.S7A are therefore composite: they provide an upper bound for the oldest admixture event, and a lower bound for the more recent one. The source of recent gene flow of Central Siberian ancestry to the Yakuts does not appear to be any of the neighboring Tungusic populations, since Yakuts completely lack the Far Eastern (red) component found in all the Tungusic groups (see below); however, given the frequency of this component in the neighboring Evenks and Evens, low levels of gene flow (1-3%) cannot be excluded. In contrast, the Far Eastern component is observed in both Dolgan subgroups, albeit at very low frequency (less than 3%). This again is consistent with historical accounts of Dolgans tracing some of their ancestry (at least 33%, based on the amount of Far Eastern ancestry) to Evenks ([82] and references therein). The substantially younger signal of European admixture in the Dolgans than the Yakuts can be explained by gene flow from Russians, known on the Taimyr from the 17th century ([82] and references therein). Indeed, analysis of the means and variance of the inferred admixture blocks in the Yakuts and Dolgans (Fig.S7B) shows that the blocks of European ancestry are longer in the Dolgans than in the Yakuts and have higher variance, thus providing corroborating evidence for additional European gene flow in Dolgans. This implies that the date of 1,470 years BP inferred for the European admixture in the Dolgans reflects both an older signal of admixture already present in the Yakuts and South Siberian populations, and more recent contact with the Russian population of the Taimyr.

In summary, our results are in good accordance with the hypothesis that the Yakuts originated in South Siberia and stem from a shared ancestral population with Buryats [75,78]. The two seemingly contradictory pieces of evidence - the absence of a Western Siberian signal in the Yakuts and the different AHGs inferred for the Yakuts (Fig.S7A and Table S2) vs Buryats (Fig.5B) – could be explained by drift and differential gene flow: on the one hand, low levels of Western Siberian ancestry could have been lost by drift in the small founding population of the Yakuts (indeed, Western Siberian ancestry is present at only 2% in the Buryats). On the other hand, recent differential gene flow after the divergence of the Yakuts (Central Siberian into the Yakuts and Asian into the Buryats), as supported by the comparatively wide blocks of Central Siberian and Asian ancestry in these populations, would have changed the configuration of their AHGs (see Fig.S8 and the simulations of the effect of additional gene flow from a population that was involved in an early admixture event described in the TextS1). The ethnogenesis of the Dolgans, who are linguistically very closely related to the Yakuts, but culturally very close to Evenks, has occurred in recent times and is known to have been influenced by the neighboring Yakuts, Evenks and Russians. Indeed, all our analyses confirm that the Dolgans are most closely related to the Yakuts and the Evenks (although some additional gene flow from Nganasan cannot be excluded with the genetic data), with evidence for recent additional gene flow from a European-like source.

## Tungusic-speaking populations

Speakers of Tungusic languages are spread from the Yenisey river in the west to the Kamchatka Peninsula in the Russian Far East and from the Taimyr Peninsula in the north to China in the south (Fig.1). The Evenks and Evens, whose languages belong to the Northern Tungusic branch, are traditionally highly mobile hunters and reindeer herders who are dispersed over vast territories of Siberia [77,83], while the linguistically closely related Oroqen (Oroqen) from northeastern China are traditionally hunters and horse herders. In contrast, the other Tungusic populations of the Amur region and northern China, such as the Hezhen and Xibo, speak languages belonging to the Southern Tungusic and Manchu branches. The Hezhen are traditionally fishers and hunters [84], while the Xibo are agriculturalists [85].

In the PCA, the Even and Evenk subgroups sampled all across Siberia show remarkably little genetic differentiation; the Evenk and Even populations are differentiated only along PC3 and PC4 (Fig.2, Fig.S6) and along PC2 in the analysis comprising only the Siberian populations (Fig.S9). Surprisingly, there is no observable structure within either of these populations, even though the samples are from locations that are up to 2700 km apart.

Analysis of the IBD blocks suggests that the Evenks and Evens have a lower effective population size than the Tungusic peoples of the south (e.g. the Oroqen), as the former share more IBD segments within the population than the latter (Fig.S3). This is also evident in the pattern of genome-wide LD, where the Oroqen have lower genome-wide LD values than the Evens or the Evenks (Fig.S4A). Overall, Evenks and Evens share a large number of long IBD segments, and they also share such segments with neighbouring populations: the Evenks with the Nganasan, Dolgans, Yukhaghirs and Nenets, the Evens with the Yukhaghirs, Dolgans and the Yakuts (Fig.S5).

The results of the ADMIXTURE analysis (Fig.1) reveal that the Tungusic populations from all across Siberia, with the exception of the Xibo, trace their ancestry to three sources broadly classified here as Central Siberian (blue), Asian (pink) and Far Eastern (red). The Central Siberian component is higher in the west than in the east and south, and ranges from 85% in the Evenk subgroups and 64% in the Evens of Kamchatka to 40% in the Oroqen, 24% in the Hezhen, and 15% in the Xibo. The Far Eastern component is highest in the easternmost Even subgroups and lowest in Tungusic populations of western and southern Siberia; this ranges in frequency from 28% in the Evens of Kamchatka to 6% in the Evenks, 5% in the Oroqen, 4% in the Hezhen, and 2% in the Xibo. The Asian component is seen at highest frequency in the southeast, where it reaches 54% in the Oroqen and 71% and 80% in the Hezhen and the Xibo, respectively, while it is present at around 10% in the Evenks and Evens. The Tungusic peoples are further characterized by the complete absence of any European-like ancestry (except for the Xibo, who have 2% of this ancestry component). Analysis of the ancestry block width inferred by PCAdmix (Fig.S10B) indicates that the blocks of Central Siberian ancestry are wider and the variance around the mean is higher in the Evenk groups, while the blocks become narrower and the variance decreases in the Oroqen, Hezhen, and Xibo. This pattern is compatible with a scenario of additional gene flow from a population of mainly Central Siberian ancestry into the western Tungusic groups (Evenks, and Evens from Sakkyryyr, Sebjan and Tompo). In addition, it has been shown previously with mtDNA and Y chromosome data that the Sakkyryyr and Sebjan Even subgroups have experienced substantial amounts of admixture from Yakuts [36,86]. Our results based on autosomal data confirm this conclusion, as the Sakkyryyr Evens are much closer to the Yakuts than to the other Tungusic groups in all measures of genetic relatedness based on IBDs, while the Sebjan and Tompo Evens appear substantially closer to the Yakuts than do the Berezovka and Kamchatka Evens (Fig.S5). Also, the ADMIXTURE results show that Sakkyryyr Evens carry around 1.5% European ancestry (presumably contributed by the Yakuts), and assuming that the elevated levels of Asian ancestry in the Sakkyryyr, Sebjan and Tompo Evens in comparison to the Kamchatka and Berezovka Evens was contributed via admixture from the Yakuts, we can roughly estimate that they have received 14-40% of Yakut gene flow (also see Fig.S1 for the ADMIXTURE results at K=7). On the other hand, the widest blocks of Asian ancestry, and the highest variance around the mean, are observed in the Oroqen, Hezhen, and Xibo (Fig.S10B). This suggests substantial recent gene flow from an Asian source population into the southern Tungusic peoples. Similarly, the distribution of the Far Eastern ancestry blocks is consistent with some additional contact between the Even subgroups closest to Kamchatka and the Koryaks, although this gene flow was not as abundant as the Central Siberian admixture detected in the Evenks and western Even subgroups or the Asian admixture into the Tungusic populations of the south (Fig.S10B).

The AHGs therefore yield a composite picture of the successive layers of differential admixture experienced by the Tungusic subgroups, leading to different configurations: ((Central Siberian, Asian) Far Eastern) for the Evenks and western Even subgroups, and ((Central Siberian, Far Eastern) Asian) for eastern Evens, Oroqen and Hezhen (Fig.S10A). Since the two eastern Even subgroups (Berezovka and Kamchatka), as well as the Oroqen and Hezhen have experienced the least amount of recent gene flow from a population of Central Siberian ancestry, we assume that their AHG reflects the ancestral admixture history of all of the Tungusic-speaking populations. Later substantial gene flow from a source or sources of predominantly Central Siberian ancestry in the Evenks and western Even subgroups (Sakkyryyr, Sebjan, and Tompo) will have considerably lowered the estimated age of Central Siberian admixture for these groups, leading to this being reconstructed as the most recent event (see Fig.S8 and the simulations of the effects of additional gene flow from a population that was involved in previous admixture described in TextS1). We therefore did not date the admixture events for the Evenks and western Even subgroups, but only show the inferred AHG (Fig.S10A). Substantial gene flow from a population of predominantly Asian ancestry will have lowered the estimated age of the Asian admixture in the Tungusic populations of the Amur region, but this would not have altered the AHG configuration.

The earliest ages estimated for the first admixture event are observed in the Oroqen (3,120 ya; CI 2,700 – 3,630) and Hezhen (3,120 ya; CI 2,910-3,930) and are likely to reflect the true age of the ancestral admixture event (as these populations are least affected by additional gene flow from a Central Siberian or Far Eastern population). Similarly, since the Berezovka and Kamchatka Evens have experienced the least amount of recent Asian admixture, the date of 1860 ya (CI: 1710-2160) can be assumed to most closely approximate the age of the ancestral admixture event involving a population of Asian ancestry. This Asian admixture event is likely to have occurred in the south, since all the Tungusic populations carry the Asian ancestry component. Thus, the expansion of the Evenks and Evens to the north and west across Siberia cannot have started earlier than ∽1,860 years ago, and could have been associated with the Xiongnu empire of Central Mongolia, as discussed in TextS2. Previous studies have suggested either Lake Baikal [87] or the Amur River (reviewed in [88]) as the place of origin of the Tungusic populations. Based on our results, we can reject the area around Lake Baikal, as all populations in our study from this area have European-like ancestry tracing back more than 3,200 years ago, and the Tungusic-speaking populations lack this substrate. We therefore suggest the Amur River as the place of origin of the Tungusic populations, which is also in good agreement with the archaeological evidence (see TextS2 for details).

### Samoyedic populations

The Samoyedic languages spoken by traditionally semi-nomadic reindeer-herding as well as hunting populations of western Siberia and the Yamal and Taimyr Peninsulas of the northernmost Russian Arctic belong to the Uralic language family, which also includes Finno-Ugric languages such as Finnish and Hungarian in Europe and Khanty in western Siberia. From historical data it is known that some Samoyedic languages were spoken in the Altai region of south Siberia too, but these are now extinct [89]. As shown by the PC analysis, the Nganasan are separated from their linguistic relatives, showing affinities with Evenks and Evens instead. The separation of the Nganasan from the other Samoyedic speakers is further underlined by the ADMIXTURE analysis and the analysis of the genome-wide LD (Fig.1 and Fig.S4C).

The distinguishing characteristic of the Samoyedic-speaking populations other than the Nganasan is that they trace their ancestry to at least three different sources. Both the Selkup and the Nenets have substantial proportions of Western Siberian (Selkup: 52%, Yamal Nenets: 48%, Taimyr Nenets: 44%) and Central Siberian (Selkup: 26%, Yamal Nenets: 44%, Taimyr Nenets: 50%) ancestry. They also have European-like ancestry (Selkup: 13%, Yamal Nenets: 4%, Taimyr Nenets: 2%). In the Nganasan we observe neither a Western Siberian nor a European ancestry component. In fact, already at K=4 individuals in this population are ascribed their own component (blue; cf. Fig.S1), indicating a different history of this population compared to the other Samoyedic peoples.

AHG analysis (Table S2) indicates that in the Selkup and Nenets the Western Siberian gene flow is the latest contribution (Fig.S11A). Furthermore, wider than average blocks of Western and Central Siberian ancestry, and higher variance around the mean, suggest that all populations have experienced multiple events of gene flow from these sources (Fig.S11B). Therefore, the dates of admixture reported below are composite dates, and reflect an older date of admixture as well as this more recent additional gene flow. We estimate the date of admixture between the European (light green) and Central Siberian (blue) ancestors to be 2,700 ya (CI 2,490 -3,120) in the Selkup, and 2,490 ya (CI 2,310 – 3,120) in the Yamal Nenets (Fig.S11A). We did not attempt to date this signal of admixture in the Taimyr Nenets subgroup, because the estimated proportion of admixture in this group is too low for our method to work reliably (see Fig.S12A). The date obtained for the more recent admixture with the Western Siberian ancestral group is 1,590 ya for the Selkup and the Yamal Nenets (CIs 1,470-1,710 and 1,470-1,860, respectively); the estimated date for the Taimyr Nenets is more recent at 1,170 ya (CI 1,020 – 1,260). In this group we also observe wider and more variable ancestry blocks (Fig.S11B), which is consistent with the gene flow either being more recent, or occurring over an extended period of time.

A further observation seems pertinent with respect to the origins and prehistory of Samoyedic populations: ADMIXTURE analysis reveals that both Selkup and Nenets have an ancestry component at a frequency of 3-4% that is also present at high frequency in the Naukan Yupik and Greenland Inuit populations. This ancestry is also evident in Altaians and Tuvans (at a similar frequency as in Selkup and Nenets), but is not seen in any other Siberian population included in this study (except the Chukchi, where our analyses suggest that this ancestry is the result of recent admixture with the neighboring Naukan Yupik; see below). This might point towards a common ancestry of the Samoyedic and southern Siberian populations (see TextS2 for more details).

All our analyses indicate that the Nganasan, the northernmost indigenous people of Siberia, are quite distinct from the other Samoyedic populations. They lack the Western Siberian and the European ancestry components, which are both characteristic of the other Samoyedic groups. Genetically, they appear to be much closer to the neighboring Tungusic speakers and the Yukaghirs than to the Selkup and even the neighboring Nenets. This leads us to conclude that the Nganasan have a different genetic history than the other Samoyedic groups and that they were linguistically, but not genetically, assimilated by the Samoyedic populations who around 1,000 years ago migrated from the south to the northern Arctic.

### Populations of the Russian Far East: Koryaks, Chukchi and Naukan Yupik

The Chukotka and Kamchatka Peninsulas were once part of ancient Beringia, a vast late Pleistocene glacial refugium for modern humans and a center of their dispersal into the New World [12]. The oldest reliably dated human remains from Beringia come from the Kamchatka Peninsula and are 13,000 years old [90]. Today, native populations of western Beringia – in particular, Chukchi, Koryaks and Naukan Yupik – speak distinct languages and differ in their social organization. The Chukchi and Koryaks, who speak languages of the Chukotka-Kamchatkan family, used to lead a nomadic lifestyle and practice inland reindeer herding in combination with coastal hunting and fishing as a mode of subsistence. The Naukan Yupik, who live in close proximity to the Chukchi, speak a Yupik language (which belongs to the Eskimo-Aleut language family extending across Alaska, the Canadian Arctic and Greenland) and traditionally engaged solely in coastal sea mammal hunting [91,92].

The PCA results reveal genetic affinities between the Koryaks, the Chukchi and the Naukan Yupik. The genetic relationship between the Chukchi and Koryaks is further confirmed by the ADMIXTURE analysis, which in addition suggests reciprocal gene flow between the Chukchi and Naukan Yupik, albeit with gene flow predominantly from the Naukan Yupik into the Chukchi.

Strikingly, both the short-range and long-range genomewide LD is substantially higher in the Naukan Yupik than in the Chukchi, who often reside in neighboring villages or even the same settlements. This would imply that in contrast to the Chukchi, the Naukan Yupik underwent an additional strong bottleneck. Overall, the populations of the Russian Far East extensively share IBD blocks among themselves as well as with the Yukaghirs and the neighbouring Even subgroups (Fig.3, Fig.S5). The Koryaks are observed to share fewer IBD blocks with the Naukan Yupik than the Chukchi do. This is probably due to admixture between the Naukan and the Chukchi, as also indicated by the ADMIXTURE analysis. Fitting a multiple regression of genetic distances on geographic distances reveals that the Naukan Yupik exhibit elevated genetic distances with all other populations (except for the neighbouring Far Eastern populations and Greenland Inuit) (Fig.S13). This pattern of generally elevated genetic distances, along with the genetic proximity to Greenland Inuit, is consistent with the viewpoint that the ancestors of the Naukan Yupik back-migrated to Siberia from the Americas [42,93,94]. In order to infer the probable homeland of the Nauakn Yupik in the Americas, we recalculated the regression coefficients, assigning to the Naukan Yupik the geographic coordinates of a range of locations in Alaska, Canada and Greenland that are home to the current Yupik and Inuit populations. The largest improvement was achieved when the location of the Naukan Yupik was assigned to the geographic area circumscribed by the south-west of the Yukon Territories, the western part of the Northern Territories and British Columbia (see Fig. S14) (Rsq = 0.613, p<0.001). This area is outside of the region where Yupik and Inuit settlements are attested; however, genetic admixture with the Chukchi will have reduced the genetic distance between the Naukan Yupik and Chukotka and thus will also reduce the geographic distance needed to provide the best match between genetics and geography. Given that from the perspective of geographic distances the territory of origin we reconstruct lies to the east of Alaska, we can conclude that the origin of the Naukan Yupik was neither in Chukotka nor in Alaska, but somewhere further east.

To estimate dates of admixture for the gene flow between the Naukan Yupik and Chukchi, the Koryaks and Greenland Inuit were taken as proxies for the parental populations. We estimated 7% of the Far Eastern component in the Naukan Yupik, and 40% Yupik contribution to the Chukchi. The estimated time of admixture for the Naukan Yupik is 1,590 ya (CI 1,470 – 1,710), and for the Chukchi 1,470 ya (CI 1,380 – 1,710), suggesting a single event of reciprocal admixture. The date of this admixture event is in good accordance with the time of the first appearance of the Neo-Eskimo culture in Siberia [95] (see TextS2 for more detail) as well as with the first appearance of cultural complexes in Alaska similar to the ones on Chukotka [96]. This date also coincides with severe climatic changes in Greenland [97], which could have incited large-scale migrations and relocation of peoples. It is also in agreement with the age of 1,000-2,000 years estimated for the expansion from Alaska and westward spread of the mtDNA haplogroup A2a, which is typical of the Eskimo populations of North America and Siberia [94].

## Conclusions

Figure 6, summarizes and synthesizes the results of all of our analyses. We find that practically all Siberian populations show evidence of extensive and deeply complicated admixture. Admixture events are often reciprocal, with many populations showing signals of multiple independent pulses of gene flow and/or of gene flow occurring over a prolonged period of time. Disentangling the ancestries from many different (often related) sources that mixed at different time points in the past, and often more than once, is a very challenging problem. To unravel this complexity we developed a new method, the Admixture History Graph (AHG), which allows one to easily determine the sequence of admixture events in a population with a complex admixture history. The approach was tested on simulated data, and we show that it is applicable to a wide variety of admixture scenarios (see TextS1). The information obtained via the AHG test can then be used to date multiple admixture events involving more than two sources of ancestry. To this end we adapted and tested with simulated data a previously published admixture dating method [45]. As with the other such methods [98,99], analysing continuous admixture remains a difficult task. Even though we can identify continuous admixture events by analysing the distribution of widths of ancestral blocks, the dates inferred for these events do not provide information on the beginning or the end of the gene flow, but show a composite of these two dates instead.

**Fig. 6.**
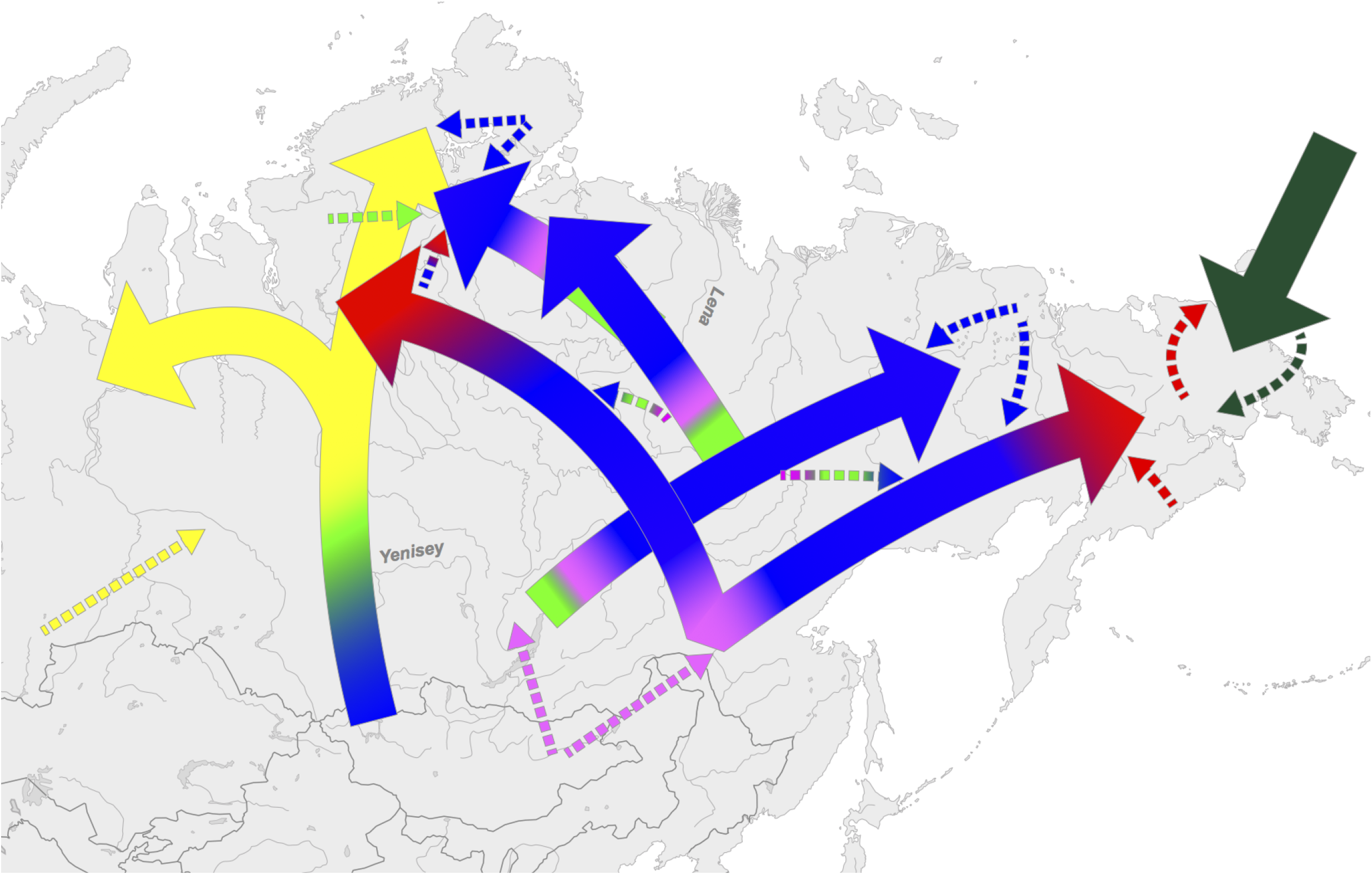
A schematic representation of the suggested migrations and directions of gene flow.

Our results indicate that the populations of Siberia share a relatively recent history, with none of the admixture dates detected being older than ∽4,500 years. This confirms previous suggestions of recent population replacements in this region [22,100,101] and fits with other evidence of depopulation of Siberia during the LGM [21]. Modern Siberian populations are therefore not expected to be good proxies for the ancestors of the populations that settled the New World.

The most extensive admixture has clearly happened in South Siberia, which is also the area to which several of the central and northern Siberian populations trace their origins. Thus, the Yakuts show clear evidence of having originated in an area near Lake Baikal and of stemming from a common ancestral population with the Buryats. The Selkup and Nenets, too, show signs of a southern origin and possibly shared ancestry with the Tuvans. In contrast, the Tungusic-speaking populations are likely to have originated not in the Lake Baikal area, as suggested previously [87], but in the Amur region [88]. Some of these recent expansions from southern to northern Siberia may have been driven by the advent of pastoralism, especially reindeer domestication, which is known in South Siberia from around 3,000 ya [56–59].

Furthermore, our results demonstrate that the European ancestry component detectable in many populations of southern, central, and northern Siberia is not the result of post-colonial Russian admixture as may have been expected [37] and as was suggested on the basis of ALDER analyses [38]. Rather, with the exception of the Dolgans and the Samoyedic-speaking groups of western Siberia, the European ancestry represents one of the most ancient components of the complex admixture history of Siberian populations.

In summary, our new methods for determining the sequence and date of multiple admixture events has enabled us to disentangle the extremely complex patterns of prehistoric admixture among Siberian populations. These methods will also be useful in further investigations of population prehistory and gene flow in other regions of the world.

## Materials and Methods

### Ethics statement

The collection of the samples was approved by the Ethics Committee of the University of Leipzig and the Ethics Committee of the Research Centre for Medical Genetics, Russian Budgetary Federal Institution, Moscow. Written consent to use their samples was obtained from all donors after explanation of the aims of the study.

### Genotypes

We genotyped 96 Siberian samples using Illumina 660W-Quad arrays. Quality filtering was performed as described previously [102]. These data were merged with Illumina genotypes from three other studies [40–42]. After quality filtering and data integration, 542 individuals and 353,357 autosomal SNPs remained for analysis. Not all samples were used for all of the analyses.

The dataset (Fig.1 and Table S1) includes four different Turkic-speaking groups: Altaians and Tuvans (South Siberia), Yakuts (three groups from the Sakha Republic), and Dolgans (from the Taimyr peninsula and the northwest of the Sakha Republic); four Mongolic-speaking populations (Buryats, Mongolians, Daurs, Tu); four Uralic populations: Samoyedic-speaking Nganasan (Taimyr peninsula), Nenets (sampled on the Taimyr and Yamal peninsulas), and Selkup, as well as the Finno-Ugric Khanty from western Siberia; and Tungusic-speakers from five ethnolinguistic groups representing all the branches of the Tungusic family: northern (Evens, Evenks and Oroqen), southern (Hezhen), and Manchu (Xibo). The Even samples come from five and the Evenk samples from two different locations across Siberia. Furthermore, we included Yukaghirs and Kets (who speak an isolate language and the last surviving language of the Yeniseic family, respectively), and populations of the Russian Far East: Chukchi, Koryaks (who speak languages of the Chukotko-Kamchatkan family) and Naukan Yupik (Eskimo-Aleut speakers). For comparative purposes we augmented the dataset with Aleuts as well as Inuit populations of Greenland, and added European (Russians, French and Tuscan and Bergamo Italians), Han Chinese, and Japanese populations from the HGDP (for detailed information on the populations and the provenance of the samples see Table S1).

#### Data Curation

Quality filtering was performed as described previously [102]. Twenty five individuals were removed before analysis: three due to low genotyping quality (one Nganasan, one Yamal Nenets and one Taimyr Nenets), one because it was identified as a potential close relative to others (Yamal Nenets), while the others were removed because they were determined via PCA to be outliers relative to their population (one Yamal Nenets, one Taimyr Nenets, one Yakut, four Nganasan, one Altaian, six Koryaks, one Selkup, one Japanese, five Chukchi).

The Yakut, Nganasan, Chukchi and Tundra Nenets populations from Reich et al. 2012 [42] (TableS1) were excluded from the IBD, regression and dating analyses because no geographical coordinates were available for these samples, and because we had other samples from the same populations (TableS1) whose PCA and ADMIXTURE results overlapped with those of the Reich et al. samples (Fig.2, Fig.1).

Prior to running the AHG calculation, we removed: a) one individual from the Selkup population, as this sample is an outlier with respect to others in terms of its admixture profile: it is the sample with the highest proportion of Central Siberian and the lowest proportion of European ancestry (45% and 4%, respectively); b) one Yamal Nenets sample, as for this sample we estimated 28% European admixture (while the population average is 4%); c) two Xibo samples with no Far Eastern ancestry (present in all other Tungusic-speakers) and higher than the population average of European ancestry (19% and 9% vs population average of 2%); d) two Taimyr Dolgans with no European ancestry, while this ancestry is present on average at 12% in the remaining Dolgans; e) in the HGDP sample of Yakuts, European admixture ranges from 4% to 30%; while in the Verkhoyansk Yakuts it ranges from 4% to 7%; therefore, to exclude recent Russian admixture we limited the analysis in the HGDP Yakuts to individuals with European admixture not exceeding 8% (this removed 11 individuals).

Furthermore, PCAdmix runs failed to confidently resolve ancestry for some chromosomes of Altaians and Tuvans (for these populations ancestry was inferred using four source populations). These chromosomes were removed prior to the admixture dating analysis. In total, 66 and 78 haploid chromosomes (the data were phased with BEAGLE, see Materials and Methods) were removed from the Altaians and Tuvans, respectively. In addition, for the same reason in the Tuvans chromosome 20 was completely removed prior to dating.

### Statistical Analyses

#### Exploratory analyses of genetic variation

Principal components analysis was performed using the StepPCO software [45] on 353,357 markers for the entire dataset and on 504,356 markers when analyzing a subset comprising only the Siberian populations.

We used the ADMIXTURE software [46] to infer individual ancestry components and admixture proportions; the LD pruning for this analysis was done using the PLINK tool [103] with the following settings: –indep-pairwise 200 25 0.4 [104], which reduced the dataset to 148,620 markers. We ran ADMIXTURE for K=3 through K=10, and performed ten independent runs for each value of K. We confirmed consistency between runs and used the cross-validation procedure implemented in ADMIXTURE to find the best value of K (Fig.S2).

#### Methods of demographic inference

For subsequent analyses, genetic distances between SNPs were interpolated using genetic coordinates and recombination rates estimated as part of the HapMap [105] and the 1000 Genomes Projects [106], and obtained using the SNAP tool [107].

Genome-wide LD calculation was performed using custom scripts, as described in [102], based on the genetic map downloaded from the Illumina webpage (http://support.illumina.com/array/array_kits/human660w-quad_dna_analysis_kit/downloads.ilmn). In particular, we binned genotype data from each population into 50 evenly spaced recombination distance categories (0.005 cM - 0.25 cM). For each population and for every pair of SNPs in each distance category we calculated the squared correlation in allele frequencies (Rsq) by selecting from each population 11 individuals showing the least amount of admixture (as determined by ADMIXTURE). For this analysis individuals from different sampling locations of the same population were grouped. To account for missing data, each pairwise measurement was adjusted by sample size.

The data were phased using BEAGLE v3.3.2. IBD segments were inferred using the fastIBD method implemented in BEAGLE [108,109]. As recommended by the developers, we ran the algorithm 10 times with different random seeds. The results were then combined and post-processed using custom scripts, but following the procedure described previously [50]. In particular, we extracted only those IBDs seen at least twice in the ten BEAGLE runs and kept only those with a significance score lower than 10^9. To remove gaps introduced artificially because of low marker density or potential switch error, we merged any two segments separated by a gap shorter than at least one of the segments and no more than 5 cM long [50]. For this calculation individuals were grouped in two ways: according to their population designation; and by sampling location. All results were adjusted for differences in sample size, and blocks shorter than 2cM were excluded [50]. To run PCA based on the results of the IBD calculation, we used the ade4 R package [110]; as a distance matrix we used the inverse of the pairwise matrix based on the number of shared IBD blocks (such that two populations sharing the smallest number of IBD blocks would be furthest apart from each other in terms of distance).

Multiple regression on genetic and geographic distance matrices was performed and the statistical significance of the regression coefficients was determined using the ‘‘ecodist” R package [111]. For this, the inverse IBD-sharing matrix was used as a measure of genetic distance between populations. Geographic coordinates for the HGDP populations were taken from Ramachandran et al 2005 [54], while the sampling locations for the samples introduced in this study were used as recorded at the time of sample collection, with Google Maps used to determine longitude and latitude for these locations. For the populations taken from other studies, where the exact sampling location was unknown, we used the coordinates of a central point of the population’s current place of residence (province or region, as listed in Table S1). Geodesic distances were calculated using the distonearth R function [55,112]. Rivers were added as an additional factor into the multiple regression model as putative facilitators of gene flow as follows: the Ob as a facilitator of gene flow between the Yamal Nenets and Khanty; the Yenisei as a facilitator of gene flow between the Taimyr Nenets, Selkup, Kets, Evenks, Tuvans, and Altaians; the Amur as a facilitator of gene flow among Hezhen, Oroqen, and Daurs; the Lena for gene flow between Buryats, Yakuts, and the Even subgroups from Tompo, Sakkyryyr and Sebjan; and finally the Kolyma as a facilitator of gene flow between Yukaghirs and the Evens from Berezovka. Two populations were assigned zero distance between them if they were connected by a river, and a distance of one if they were not.

#### Analyzing and dating multiple admixture events

To analyze signals of admixture in the Siberian populations involving more than two ancestral populations we adopted the following workflow: the data were phased with the BEAGLE software [108], then the PCAdmix tool was used to infer ancestry along individual chromosomes [113]. Unlike other existing methods for ancestry estimation, this algorithm allows for simultaneous inference of admixture from more than two sources of ancestry. In order to determine the sequence of admixture events we developed a new approach, the Admixture History Graph (AHG, see below for details), based on the results of the ADMIXTURE analysis. Finally we applied our previously published wavelet transform analysis [45], adapted to include complex admixture scenarios, to the results obtained from PCAdmix and using the information on the sequence of events obtained from the AHG, to estimate the dates of the different admixture events (see TextS1 for validation). To aid interpretation, we also analyzed the length distribution of blocks of different ancestry, as inferred by PCAdmix.

##### PCAdmix

Our choice of parental populations for the PCAdmix analysis was guided by the results of the ADMIXTURE analysis for K=6, and populations that had been assigned their own ancestry component at this value of K were chosen as proxies for the true ancestral populations. Specifically, for the European ancestry component we used Italians as the parental group, for the Western Siberian component we used the Khanty, for the Central Siberian component we used the Nganasan, for the Asian component we used the Han Chinese, for the Far Eastern component we used the Koryaks, and for the Eskimo component we used the Greenland Inuit. We assigned an equal number of individuals per parental group and selected only those individuals with not more than 1% admixture. Since Greenland Inuit, whom we used as a proxy parental population for the admixture signal observed in Chukchi and Naukan Yupik, are themselves heavily admixed, we made an exception for this group and accepted up to 10% admixture in this parental population (as only three individuals met our 1% admixture criteria). Since wavelet transform analysis (see below) requires 2^n windows per chromosome [45], we used custom scripts when running PCAdmix to ensure conformity to this requirement while maintaining relatively equal numbers of SNPs per window regardless of chromosome size.

##### Admixture History Graph (AHG)

To disentangle the admixture chronology in populations with complex admixture histories, we developed a new approach referred to here as the Admixture History Graph (AHG). The approach is based on the following idea: if we consider an admixed population with only two sources of ancestry from parental populations A and B, the contributions of A and B among the individuals in the population vary. When a second admixture event occurs, bringing into the population new ancestry C, the proportion of ancestry A and B in each individual becomes adjusted by an amount which depends on the proportion of the component C in the individual (Fig.7A). This means that the distribution of the first two ancestry components A and B across individuals in the new population will covary with C, yet the ratio of A and B throughout the population will be independent from C. Thus the covariance of the recent ancestry C and the ratio of the two older ancestries A and B should be zero. To determine the sequence of admixture events we can therefore simply compare all possible orderings of the three ancestries A, B, and C and choose the one which produces the least absolute value of covariance. The same reasoning applies for populations that have experienced more than two admixture episodes (Fig.7B). In such situations, we test the configuration of all possible trios of A,B,C and D, and then reconstruct the full graph based on the orderings of likely configurations.

**Fig. 7.**
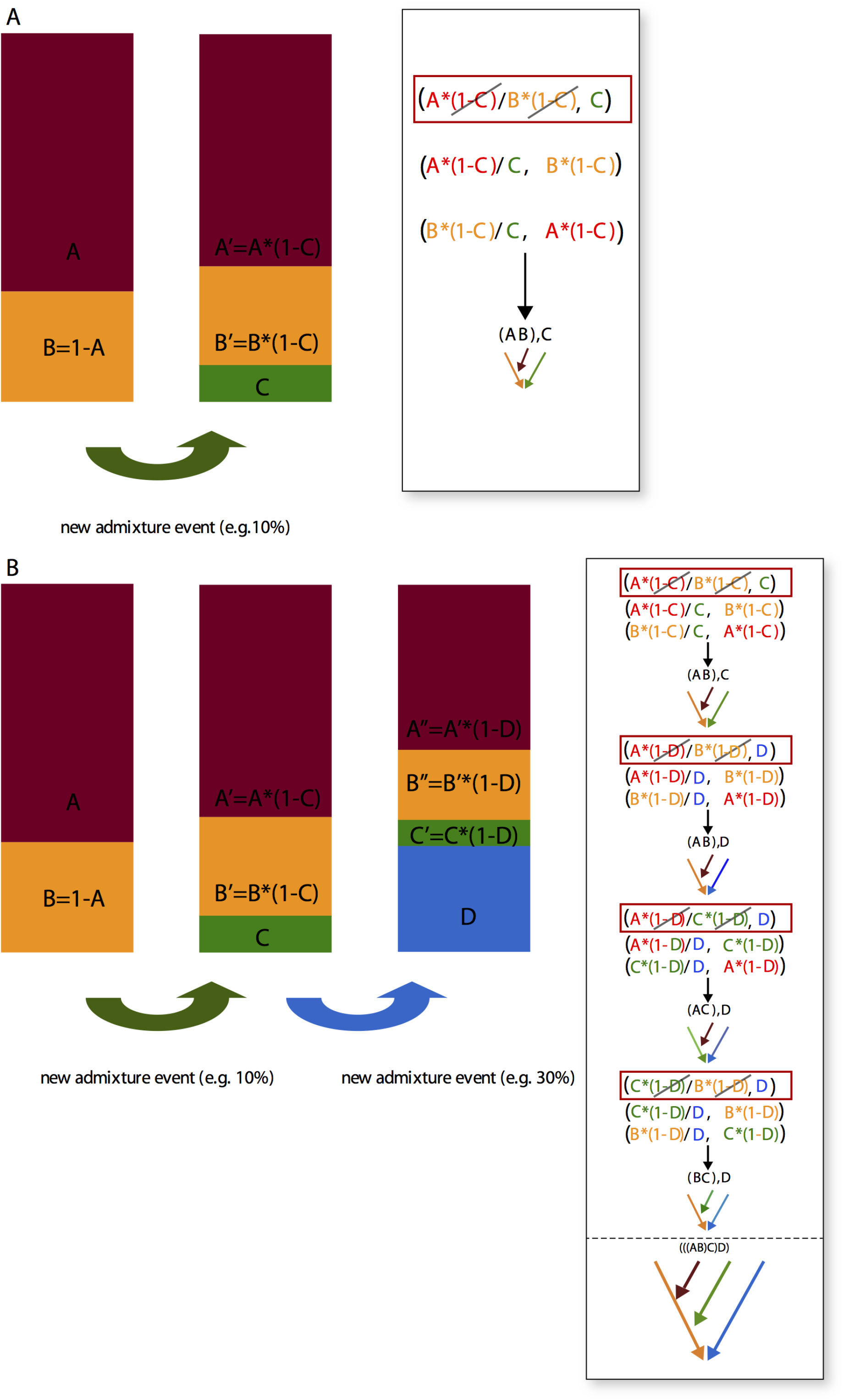
Schematic representation of the AHG approach. (A) When an already admixed population with ancestry coming from two different sources A and B experiences another episode of admixture, which brings into this population a new ancestry component C, the proportion of ancestry A and B in each individual in this population becomes adjusted by an amount which depends on the proportion of the component C in the individual. While before the admixture episode the proportion of A and B among the individuals in the population varied independently of each other, after the new admixture episode A and B start to co-vary because due to the influx of C each of them has been adjusted by the same amount. The test is based on the detection of this co-variance between A and B, as is further illustrated in the inset. (B) An example involving additional gene flow from ancestry D.

The performance of this new approach was tested with simulated data. In general, the order of admixture events was identified correctly by the algorithm and with great precision for a wide variety of scenarios and even for admixture events involving four sources of ancestry. The performance of the test is not affected by small sample sizes. However, when sample sizes are small (N=10), the variance in the sample starts to influence the results: namely if the variance around the mean of all ancestry components in the sample is identical, it obscures any existing covariance between them, and the test becomes unreliable. In addition, if the final admixture proportion of one or more components in the population under study is very low (below 5%), the accuracy of the method decreases (see TextS1 for details).

##### Wavelet Transform Analysis

Once the sequence of admixture events was determined by the AHG approach, we first considered the most recent admixture episode. PCAdmix estimates ancestry along individual chromosomes and assigns chunks of the genome to any of the three or more ancestral populations. In estimating the date for the most recent episode, we treated the population receiving the gene flow as already admixed, and its ancestry composed of ancestry blocks originating from previous admixture events, i.e. we treated all blocks of ancestry assigned by PCAdmix to two (or more) older admixture contributors as the same, and considered this part of the genome as coming from population 1. The part of the genome assigned by PCAdmix to the most recent admixture contributor (as determined by the AHG) was considered as originating from population 2. We then proceeded to use WT analysis [45] to estimate the width of ancestry blocks contributed by these two ancestral groups, and estimated the date of admixture based on simulations described previously [45], using simulations with the rate of admixture parameter closest to the populations being analyzed. To estimate the earlier admixture dates the approach was as follows: the blocks of most recent ancestry were masked, i.e. excluded from the analysis, and the signal of admixture (the rate of admixture and the size of ancestry blocks) in the remaining part of the genome was analyzed as before. The resulting reduction in genome size was taken into consideration when the result was compared to the simulated data to obtain the date of admixture estimate. Since the first version of the WT admixture dating method [45] is restricted to admixture events involving two parental populations, the adaptation introduced here to this previously published approach was validated in various ways (see TextS1 for details); these confirm that the method results in the accurate estimation of dates of multiple admixture events.

## Acknowledgments

IN acknowledges financial support from the Russian Foundation for the Humanities (14-04-00346), and BP is grateful to the LABEX ASLAN (ANR-10-LABX-0081) of Université de Lyon for its financial support within the program “Investissements d’Avenir” (ANR-11-IDEX-0007) of the French government operated by the National Research Agency (ANR). This research was funded by the Max Planck Society.

## Author Contributions

IP, RM, BP and MS conceived and designed the experiments. IP performed the experiments. IP analyzed the data. IP and RM contributed analytical tools. VS, SM, IN and VO contributed data. IP, BP and MS wrote the paper.

